# Auxin regulates endosperm cellularization in *Arabidopsis*

**DOI:** 10.1101/239301

**Authors:** Duarte D. Figueiredo, Rita A. Batista, Claudia Kohler

## Abstract

The endosperm is an ephemeral tissue that nourishes the developing embryo, similar to the placenta in mammals. In most angiosperms endosperm development starts as a syncytium, where nuclear divisions are not followed by cytokinesis. The timing of endosperm cellularization largely varies between species and the event triggering this transition remains unknown. Here we show that increased auxin biosynthesis in the endosperm prevents its cellularization, leading to seed arrest. Auxin-overproducing seeds phenocopy paternal-excess triploid seeds derived from hybridizations of diploid maternal plants with tetraploid fathers. Concurrently, auxin-related genes are strongly overexpressed in triploid seeds, correlating with increased auxin activity. Reducing auxin biosynthesis and signaling reestablishes endosperm cellularization in triploid seeds and restores their viability, highlighting a causal role of increased auxin in preventing endosperm cellularization. We propose that auxin determines the time of endosperm cellularization and thereby uncovered a central role of auxin in establishing hybridization barriers in plants.

## Introduction

In flowering plants seed development is initiated by the fertilization of two maternal gametes, egg cell and central cell, by two paternal sperm cells (1). This double fertilization event originates two fertilization products: the embryo, which will form a new plant, and the endosperm, a nourishing tissue that ensures adequate nutrient transfer from the mother plant to the developing embryo (2). The endosperm of most angiosperms is a triploid tissue, derived after fertilization of the diploid central cell. It thus contains two maternal and one paternal (2M:1P) genome copies. In *Arabidopsis*, like in most angiosperms, the endosperm initially develops as a syncytium, where nuclear divisions are not followed by cytokinesis (3). After a defined number of nuclear divisions the endosperm cellularizes (4); however, the pathways regulating this transition remain unknown. The balance of 2M:1P genome copies in the endosperm is crucial for reproductive success. Deviation from this ratio in response to hybridizations of plants that differ in ploidy frequently leads to unviable seeds, a phenomenon referred to as triploid block (5–9). Importantly, interploidy hybridizations affect endosperm cellularization: while maternal excess crosses (4x × 2x; by convention the maternal parent is always mentioned first) shift the cellularization to earlier timepoints, paternal excess hybridization (2x × 4x) cause a delay or complete failure of endosperm cellularization (8, 10). In *Arabidopsis*, the triploid (3x) embryos resulting from 2x × 4x crosses are viable and produce healthy plants when transferred to nutritive medium, revealing that failure of endosperm cellularization impairs embryo viability (10, 11). Mutations in the paternally-expressed imprinted genes (PEGs) *ADMETOS (ADM), SUVH7, PEG2*, and *PEG9* restore endosperm cellularization and viability of paternal excess 3x seeds (12–14).

In this study we show that auxin activity is strongly increased in paternal-excess 3x seeds and that the 3x seed phenotype can be phenocopied by over-production of auxin in the endosperm of diploid seeds. Furthermore, we show that down-regulating auxin biosynthesis or signalling can partly restore 3x seed viability. Overall, our data link auxin activity with endosperm cellularization and show that increased auxin activity in the endosperm establishes a post-zygotic hybridization barrier in *Arabidopsis*.

## Results

### Paternal-excess crosses lead to increased auxin activity after fertilization

Triploid seed abortion in paternal-excess (2x × 4x) crosses is characterized by the over-proliferation of the endosperm, which fails to cellularize (8), but the molecular mechanisms that lead to this phenotype are yet to be elucidated. To search for pathways potentially involved in 3x seed abortion, we compared gene expression data of WT seeds at 6 days after pollination (6 DAP) with that of WT maternal plants pollinated with pollen of *omission of second division 1* (*osd1*) (15). Mutants for *osd1* form unreduced diploid gametes (2n) and therefore can be used to mimic paternal-excess crosses when used as a pollen donor to a WT mother (12, 15). We found genes involved in auxin homeostasis to be significantly enriched among those genes that were upregulated in 3x seeds (Table 1). In particular, genes involved in auxin biosynthesis (*TAA/TAR* and *YUCCA* (16, 17)), auxin transport (PIN and PGP-type (18, 19)), and Auxin Response Factors (ARFs (20, 21)) were highly upregulated in 3x seeds, when compared to 2x seeds (Fig. 1A and Fig. 1-S1). Consistent with the transcriptome data, we found a marked increase in the activity of (22)the auxin sensor *DR5v2::VENUS* (23) in 3x seeds that was most prominent in the seed coat, suggesting that increased auxin generated in the fertilization products in response to *osd1* pollination is rapidly transported to the seed coat (Fig. 1D,E), in line with previous reports (24). Indeed, genes coding for auxin biosynthesis, as well as auxin signalling, are strongly up-regulated in the endosperm of 3x seeds compared to that of 2x seeds (Fig. 1-S2).These observations indicate that paternal-excess crosses induce increased auxin production and signalling in the endosperm of 3x seeds.

**Figure 1.**
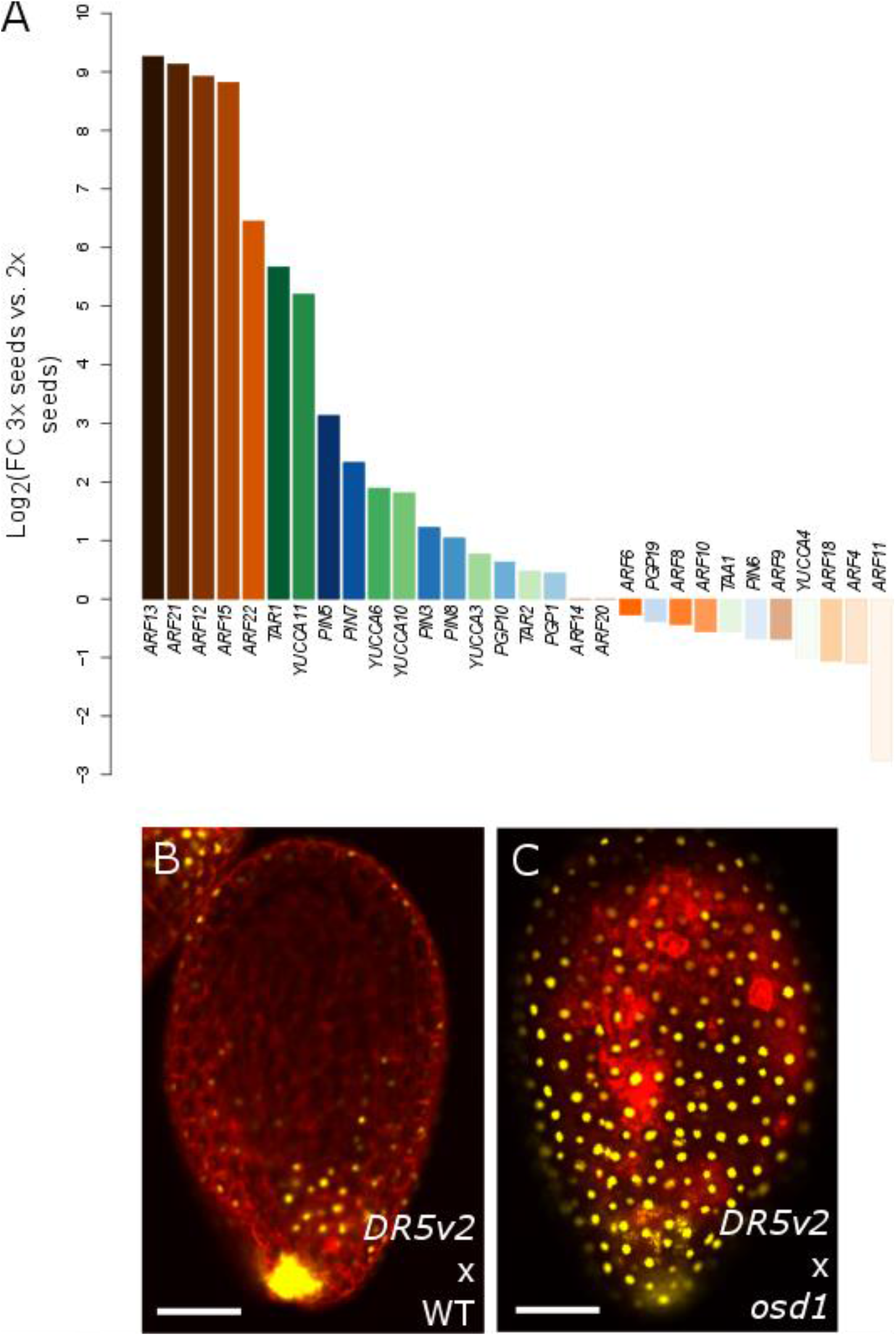
Auxin is overproduced in 3x seeds. (A) Log2-fold expression change between 3x and 2x seeds of genes coding for auxin biosynthesis (green bars), signalling (orange bars) and transport proteins (blue bars). (B-E) Auxin activity as measured by expression of *DR5v2::VENUS* in 2x (B) and 3x (C) seeds at 5 days after pollination. Pictures show representative seeds of three independent siliques per cross. Red staining is propidium iodide. Scale bars indicate 100 μm.

### Over-production of auxin in the endosperm phenocopies paternal-excess triploid seeds

Based on the finding that auxin activity is increased in 3x seeds, we addressed the question whether over-production of auxin is responsible for the endosperm developmental defects leading to 3x seed abortion. To test this hypothesis, we raised transgenic plants over-expressing the bacterial auxin biosynthesis gene *Indole Acetimide Hydrolase (IaaH)* under the control of the early-endosperm specific promoter *DD25* (25, 26). The production of the active auxin Indole 3-Acetic Acid (IAA) by IaaH relies on the availability of Indole 3-Acetamide (IAM), which was previously shown to be present in *Arabidopsis* (27, 28). Furthermore, genes coding for IAM-synthetizing enzymes are strongly expressed in the endosperm (Fig. 2-S1). Strikingly, out of 31 transgenic lines expressing *DD25::IaaH*, all showed aborting seeds that closely resembled paternal-excess 3x seeds by their dark and shriveled appearance (Fig. 2A-C). In seven lines that were analyzed in detail we found that the frequency of either partially or fully collapsed seeds ranged between 10 to 40%, which largely corresponded with the rate of non-germinating seeds (Fig. 2-S2). Embryos of *DD25::IaaH* expressing lines were retarded in growth, similar to 3x embryos (Fig. 2 D-L). In both 3x seeds and those expressing *DD25::IaaH*, embryo development progressed up to the early heart stage without noticeable differences compared to 2x WT seeds (5 DAP time-point; Fig. 2). However, from 6 DAP onwards, the embryos of 3x and *DD25::IaaH* transgenic seeds were delayed in development and did not progress beyond the torpedo stage (Fig. 2 D-L).

**Figure 2.**
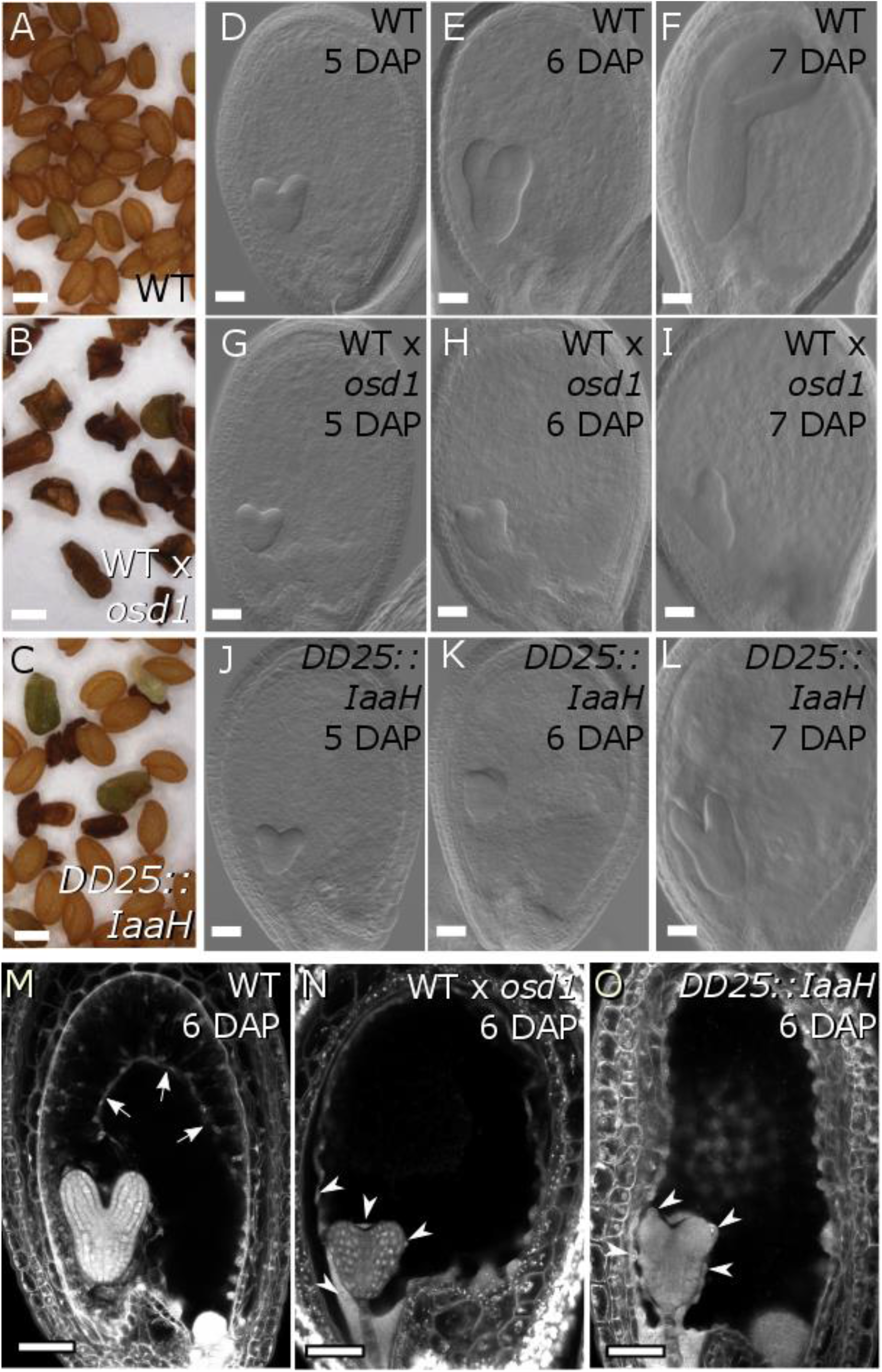
Increased auxin in the endosperm prevents cellularization. (A-C) Dry seed morphology of WT 2x (A), WT 3x (B) and *DD25::IaaH* 2x seeds (C). Scale bars indicate 0.5 mm. (D-L) Clearings of WT 2x (D-F), WT 3x (G-I) and *DD25::IaaH* 2x seeds (J-L), from 5 to 7 days after pollination (DAP). Pictures show representative seeds of three independent siliques per cross. Scale bars indicate 50 μm. (M-O) Endosperm cellularization as determined by Feulgen staining at 6 DAP for 2x seeds (M), 3x seeds (N) and 2x seeds expressing *DD25::IaaH* (O). Pictures show representative seeds of 10 independent siliques per cross. Arrows indicate cellularized peripheral endosperm and arrowheads indicate free endosperm nuclei surrounding the embryo. Scale bars indicate 50 μm. WT, wild type.

The endosperm of seeds derived from paternal-excess crosses fails to cellularize (8); therefore, we tested whether seeds expressing *DD25::IaaH* showed a similar developmental defect (Fig. 2M-O and Fig. 2-S4). Endosperm cellularization of 2x WT seeds initiated around 5 DAP and was almost complete at 7 DAP (Fig. 2-S4). Consistent with previous reports (8), in 3x seeds derived from paternal-excess crosses the endosperm failed to cellularize and free endosperm nuclei could be seen surrounding the embryo. Importantly, *DD25::IaaH* expression induced a similar phenotype and many seeds showed no signs of endosperm cellularization even at 7 DAP (Fig. 2-S4). These observations indicate that over-production of auxin in the endosperm is sufficient to impair its cellularization.

To test whether the phenotypes observed in *DD25::IaaH* lines are indeed caused by over-production of auxin in the endosperm, we crossed WT plants with pollen from *DD25::IaaH* plants. Indeed, we observed the same seed phenotypes when the transgene was inherited through pollen, confirming that endosperm-produced auxin is causal to this phenotype, and ruling out that the effect originates in maternal sporophytic tissues (Fig. 2-S2). Furthermore, when crossing maternal plants expressing the auxin reporter *DR5v2::VENUS* with pollen carrying the *DD25::IaaH* transgene, we observed a significant increase in VENUS fluorescence, similar to what is observed in paternal-excess crosses (Fig. 2-S2 and Fig. 1D-E). Together we conclude that increased auxin production in the endosperm prevents endosperm cellularization, leading to a phenocopy of paternal-excess 3x seeds.

### *PEGs* and *AGAMOUS-LIKE* genes are only partly de-regulated by auxin over-production

Triploid paternal-excess seeds are characterized by a strong deregulation of PEGs and genes coding for AGAMOUS-LIKE (AGL) MADS-box transcription factors (12, 29). We tested whether overproduction of auxin causes a transcriptional phenocopy of 3x paternal-excess seeds by analyzing expression of *PEGs* and *AGLs* that were previously shown to be strongly deregulated in 3x seeds (13, 30). While PEG genes *ADM* and *PEG9* were not significantly deregulated in seeds of *DD25::IaaH* expressing plants compared to 2x WT seeds, *PEG2* and the *AGL* genes *PHE1, AGL62*, and *AGL36* were expressed at significantly higher level in auxin-overproducing seeds (Fig. 3). However, their level of deregulation remained substantially lower compared to 3x seeds. This data suggest that auxin acts either independently of the pathways previously shown to affect 3x seed abortion (13, 30, 31) or, alternatively, that auxin signalling is downstream of PEG and AGL functions in the endosperm.

**Figure 3.**
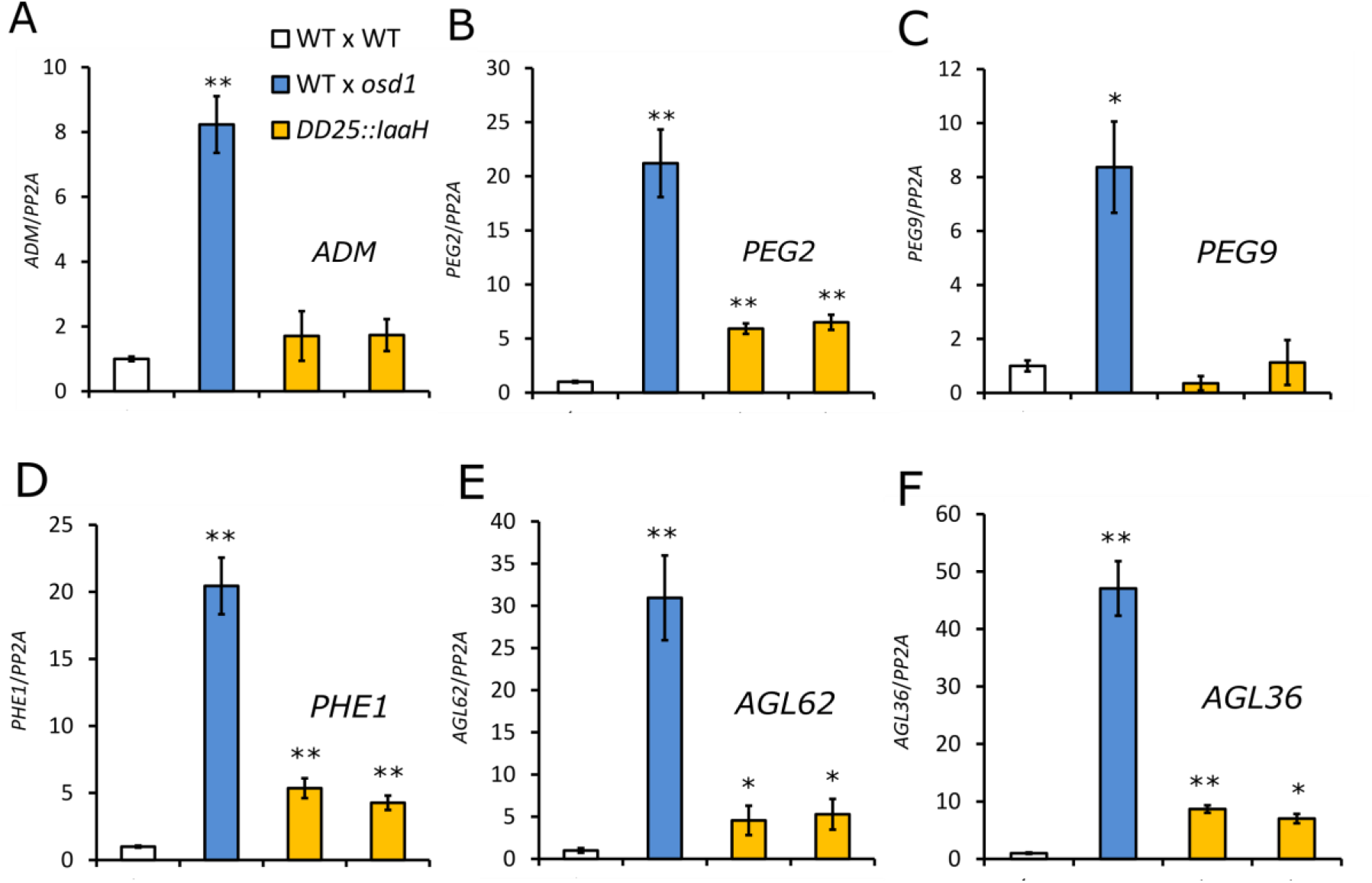
*PEGs* and *AGL* genes are not substantially deregulated in *DD25::IaaH* transgenic seeds. Relative gene expression in seeds at 6 days after pollination (DAP), as determined by RT-qPCR, in 2x WT, 3x WT and 2x *DD25::IaaH* transgenic seeds of two independent lines for *ADM* (A), *PEG2* (B), *PEG9* (C), *PHE1* (D), *AGL62* (E) and *AGL36* (F). Results from a representative biological replicate are shown. Three technical replicates were performed and error bars indicate standard deviation. Differences are significant for Student’s T-test for p<0.05 (*) or p<0.001 (**). WT, wild-type.

### Decreased auxin biosynthesis and signalling suppress triploid seed abortion

To address the question whether endosperm failure in 3x paternal excess seeds is due to over-production of auxin, we analyzed whether mutants for either auxin biosynthesis (*wei8 tar1 tar2*−1/+) (17), or auxin signalling (*axr1*) (32) could suppress 3x seed abortion. We generated 4x WT and *wei8 tar1 tar2/+* plants by colchicine treatment and used these plants as pollen donor in crosses with 2x WT or *wei8 tar1 tar2/+* mutant maternal plants. In the 2x × 4x WT cross, around 70% of the seeds were fully collapsed (Fig. 4A). In contrast, only 20% of 3x *wei8 tar1 tar2/+* seeds were fully collapsed and the germination rate of mutant 3x seeds was nearly doubled compared to WT (Fig. 4B), revealing that decreased auxin biosynthesis can suppress triploid seed abortion. To substantiate these findings we tested the effect of the auxin signalling mutant *axr1* in suppressing 3x seed abortion. Using the *osd1* mutant as pollen donor resulted in around 50% fully collapsed 3x seeds, while only 20% of 3x seeds were fully collapsed when using the *axr1 osd1* double mutant as pollen donor (Fig. 4C). The *axr1* 3x seeds were phenotypically distinct from 2x WT seeds by having a box-shaped phenotype (Fig. 4-S1); however, many of these seeds were viable and germinated at a rate of 40%, compared to 9% of 3x WT seeds (Fig. 4D-F). These results demonstrate that decreased auxin signalling can suppress paternal-excess seed abortion. The fact that 2x *axr1* seeds are largely viable, compared to only 80% viability of 2x *wei8 tar1 tar2/+* seeds (Fig. 4-S1), likely accounts for the decreased viability of 3x *wei8 tar1 tar2/+* seeds when compared to 3x *axr1* (Fig. 4B,D).

**Figure 4.**
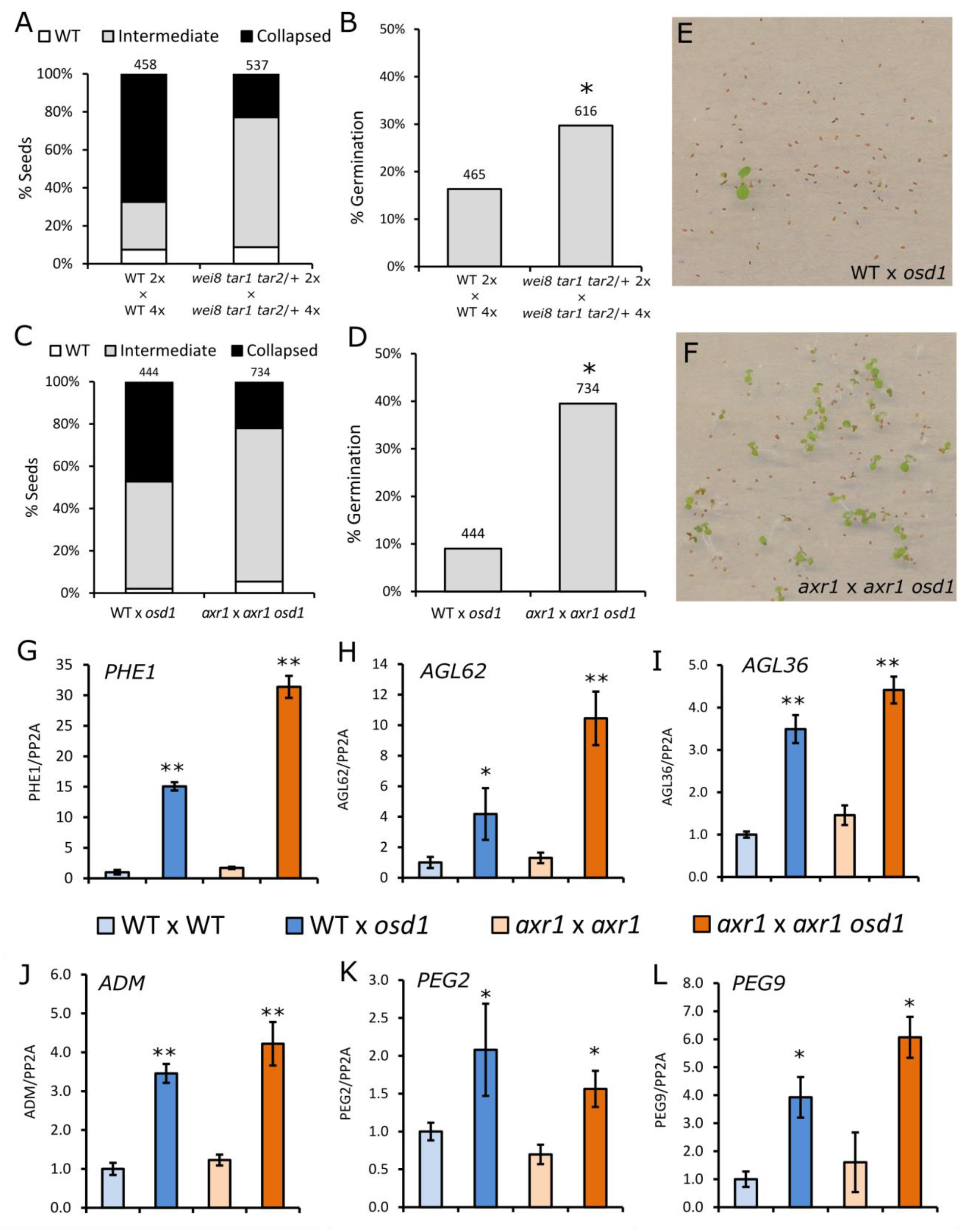
Mutants in auxin biosynthesis and signaling suppress triploid seed abortion. (A, B) Phenotypic classification of 3x seeds in WT background and in the *wei8 tar1 tar2* auxin biosynthesis mutant (A), and their corresponding germination rate (B). (C, D) Same as for (A) and (B), but for the auxin signalling mutant *axr1*. Seed classification was done according to Fig. 2 – S2. Numbers on top indicate number of seeds assayed. Differences between WT and mutant seed germination in (B) and (D) are significant for Chi-square test for p<0.0001 (*). (E, F) Representative image of germinating triploid seedlings in WT (E) and *axr1* (F). (G-L) Relative gene expression in 6 DAP seeds, as determined by RT-qPCR, in 2x and 3x seeds in WT and *axr1* mutant backgrounds, for *PHE1* (G), *AGL62* (H), *AGL36* (I), *ADM* (J), *PEG2* (K) and *PEG9* (L). Results of a representative biological replicate are shown. Three technical replicates were performed and error bars indicate standard deviation. Differences between 3x seeds and each respective 2x control are significant for Student’s T-test for p<0.05 (*) or p<0.001(**).

Given that triploid seed abortion is characterized by a failure of the endosperm to cellularize, we thus tested if this process was restored in the *axr1* mutant background (Fig. 4-S2). Endosperm cellularization dynamics in 2x *axr1* seeds was similar to that of 2x WT seeds (Fig. 2 and Fig. 4-S2) and the endosperm was almost fully cellularized at 7 DAP. Although cellularization in 3x *axr1* seeds was delayed compared to 2x seeds, signs of endosperm cellularization in this mutant were clearly visible at 7 DAP, as opposed to 3x WT seeds. We thus conclude that rescue of the 3x seed abortion by reduced auxin signalling occurs by restoration of endosperm cellularization.

As discussed above, *AGL* genes and *PEGs* are strongly upregulated in 3x seeds (Fig. 3). We addressed the question whether the rescue of 3x seeds by reduced auxin signalling restored gene expression to WT levels. Thus, we tested the expression of *AGL* genes and *PEGs* in 2x and 3x WT and *axr1* seeds. For all genes tested, their expression remained significantly increased in 3x *axr1* seeds, and, with the exception of *PEG2*, was even higher in 3x *axr1* seeds than 3x WT seeds (Fig. 4 G-L). This data reveals that 3x seed rescue by decreased auxin signalling occurs independently of PEGs and AGLs and strongly suggests that auxin acts downstream of those pathways during endosperm development.

### Downregulation of auxin biosynthesis and signaling genes coincides with endosperm cellularization

The observation that over-production of auxin prevents endosperm cellularization suggests that auxin levels have to fall below a certain threshold in order for the endosperm to cellularize. To test this hypothesis we analyzed the expression of auxin biosynthesis, transport, and signaling genes in the micropylar and chalazal domains of the endosperm. Endosperm cellularization is initiated in the micropylar domain of the endosperm at around heart stage of embryo development, while cellularization in the chalazal domain occurs later when the embryo has reached the torpedo stage (33). Indeed, the expression level of the *PEGs YUC10* and *TAR1* was significantly lower in the micropylar endosperm of heart stage 2x embryos, when compared to earlier timepoints (Fig. 5A and Fig. 5-S1), consistent with the initiation of cellularization in 2x seeds. The same expression pattern was observed for genes coding for YUC11, for PGP and PIN-type transporters and for several ARFs (Fig. 5A and Fig. 5-S1). This was further confirmed by reduced activity of the fluorescent markers *YUC10::YUC10:GFP* (34) and *PGP10::GFP* (24) at the timing of cellularization (Fig. 5-S1). Downregulation of auxin-related genes was substantially less pronounced in the chalazal endosperm domain, correlating with its delayed cellularization (Fig. 5A). Furthermore, in 3x seeds, where the endosperm fails to cellularize, there was a strong increase in expression of several auxin-related genes (Fig. 5B). Coinciding with rescued endosperm cellularization in 3x *adm* seeds (12), expression of auxin-related genes became downregulated (Fig. 5B). Together we conclude that increased auxin activity prevents endosperm cellularization, revealing a central regulatory role of auxin in the transition from the syncytial to the cellularized endosperm state.

**Figure 5.**
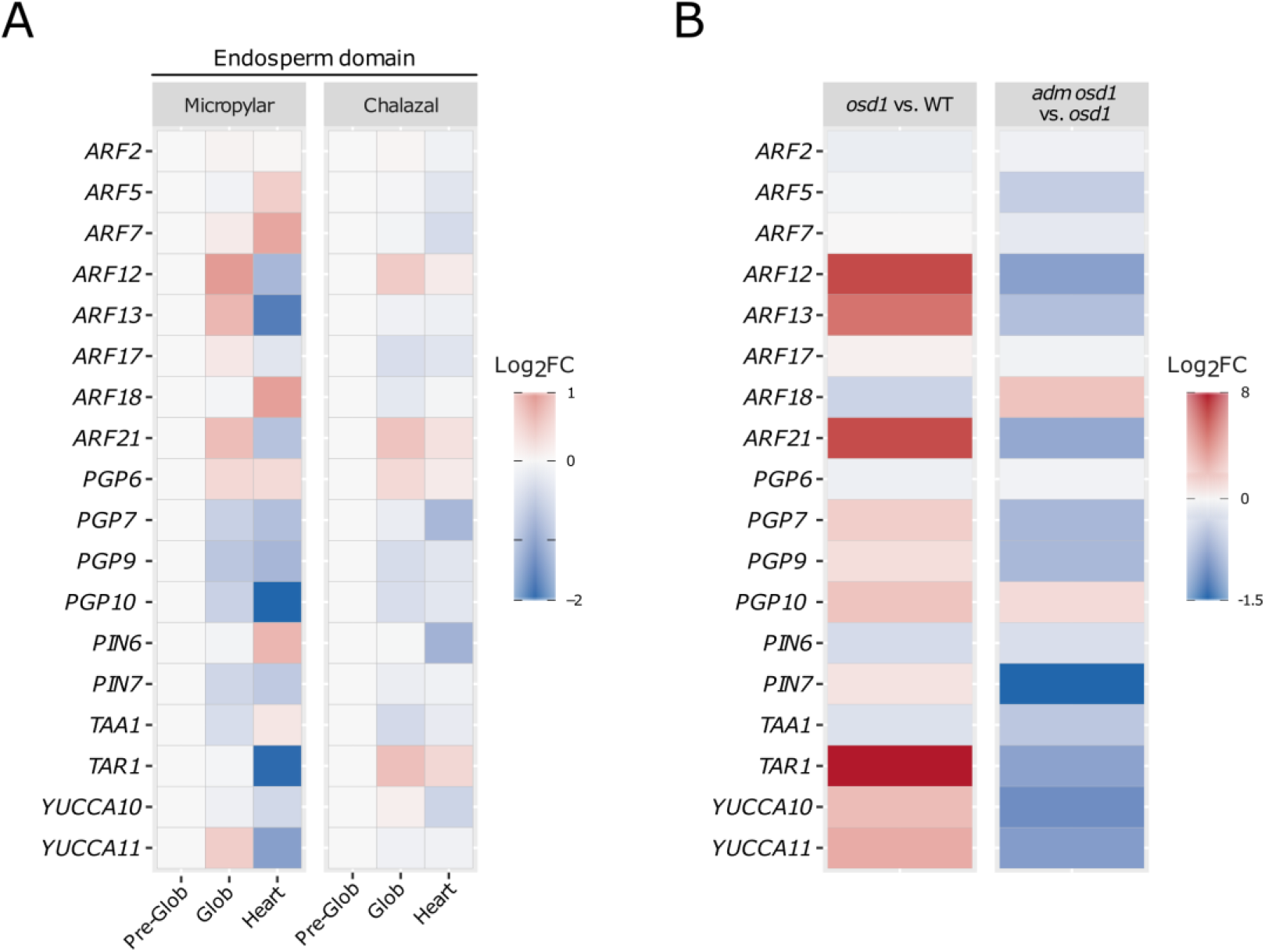
Endosperm cellularization is associated with downregulation of auxin-related gene expression. (A) Expression of auxin-related genes in the micropylar and chalazal endosperm domains, throughout different stages of seed development. Expression in each domain is normalized to the pre-globular stage and expressed as a log2-fold change relative to that stage. (B) Log2-fold change of expression of auxin-related genes in 3x (*osd1*) versus 2x seeds and 3x *adm* seeds (*osd1 adm*) versus 3x seeds (*osd1*).

## Discussion

In many plant species crosses between individuals of different ploidies have long been known to result in abortion of the progeny due to failure of endosperm cellularization, a critical process in seed development (8–10). Nevertheless, the molecular mechanisms underlying this developmental transition have remained elusive. Here we demonstrate that increased production of the plant hormone auxin prevents endosperm cellularization in 3x seeds of *Arabidopsis thaliana*. Thus, in addition to its known role to initiate endosperm development and seed coat formation (24, 35), we propose that auxin levels need to be tightly controlled at later stages of seed development to allow the endosperm to cellularize. This hypothesis is strongly supported by our findings that over-production of auxin prevents endosperm cellularization in 2x seeds and that down-regulation of auxin activity in 3x seeds restores cellularization and, consequently, seed viability. Importantly, the auxin-induced endosperm phenotype is characteristic of paternal-excess crosses, leading to uncellularized inviable seeds (8, 9). Auxin biosynthesis genes *YUC10* and *TAR1* are imprinted and paternally-expressed in the endosperm (35–37). Like many other PEGs, *YUC10* and *TAR1* are upregulated in the endosperm of 3x seeds (13), likely causing increased auxin biosynthesis. The observed strong increase of ARF expression may be a consequence of a positive feedback loop, similar to the self-sustained activation of the ARF *MONOPTEROS* during early embryogenesis (38). ARFs are transcription factors that regulate the expression of auxin-responsive genes (21, 39) and thus are able to amplify the response to increased auxin levels in the endosperm.

Our data suggests that increased auxin activity in the endosperm is likely downstream or independent of AGLs and the known suppressors of 3x seed abortion ADM, PEG2, and PEG9 (12, 13). This conclusion is based on the fact that increased auxin could induce a 3x seed-like phenotype without causing increased suppressor gene expression. Furthermore, reduced auxin signaling in *axr1* could suppress 3x seed abortion despite high expression levels of *ADM*, *PEG2*, and *PEG9*. Consistent with auxin acting downstream of ADM, most auxin-related genes being upregulated in 3x seeds became repressed in 3x *adm* seeds. We propose that endosperm cellularization can only take place when auxin levels are below a certain threshold. If this threshold is not reached, like in 3x seeds, the endosperm will fail to cellularize and the seed aborts. Interestingly, endosperm cellularization in maize occurs at around 3 DAP, clearly before the rise of auxin levels at around 9 DAP (40). The rise in auxin levels coincides with the onset of endoreduplication and cellular differentiation in the endosperm, while proliferation rates decrease. It therefore seems unlikely that endosperm cellularization failure in 3x seeds is a consequence of auxin-induced nuclear overproliferation, but that nuclear proliferation and endosperm cellularization are mechanistically unlinked. This is consistent with data showing that both processes can be uncoupled in response to interspecies hybridization in rice (41). Auxin is well known to induce changes in cell wall mechanical properties and cell wall synthesis. Auxin-induced organ outgrowth requires demethylesterification of pectin, which causes cell wall loosening (42). Auxin could have a similar role in the endosperm and by inducing demethylesterification and pectin degradation inhibit endosperm cellularization. We recently found increased demethylesterification activity in the endosperm of 3x seeds, adding support to this idea (13).

In conclusion, we have shown that auxin regulates endosperm cellularization in *Arabidopsis*. Increased auxin levels in 3x seeds negatively interfere with endosperm cellularization, uncovering a central role of auxin in establishing hybridization barriers by changing the timing of endosperm cellularization.

## Materials and Methods

### Plant material, growth conditions and treatments

The *Arabidopsis thaliana* mutant and reporter lines used were described previously: *wei8-1/-tar1/-tar2-1/+* and *wei8-1/-tar1/-tar2-2/+* (17), *axr1-12/+* (32), *osd1-1* (15), *osd1-3* (43).

Seeds were sterilized in 5% commercial bleach and 0.01% Tween-20 for 10 min and washed three times in sterile ddH2O. Sterile seeds were plated on ½ MS-medium (0.43% MS-salts, 0.8% Bacto Agar, 0.19% MES hydrate and 1% Sucrose; when necessary, the medium was supplemented with the appropriate antibiotics) and stratified at 4 °C in the dark for 48 h. Plates were then transferred to a growth chamber (16 h light / 8 h dark; 110 μmol.s^−1^.m^−2^; 21°C; 70% humidity). After 10 days seedlings were transferred to soil and grown in a growth chamber (16 h light / 8 h dark; 110 μmol.s^−1^.m^−2^; 21°C; 70% humidity).

Tetraploid plants were generated by treating two-week old seedlings with 7 μL of 0.25% colchicine. Treated plants were grown to maturity and scored for alterations in pollen size. Seeds of plants showing enlarged pollen grains were collected and the ploidy of the subsequent generation was determined in a Cyflow Ploidy Analyzer, using the Cystain UV Precise P kit (Sysmex).

### Transcriptome analysis

Analysis of de-regulated genes in triploid seeds was done using previously published RNAseq data and following previously published procedures (44). We generated a list of overexpressed genes in 3x seeds by filtering all genes with Log2FC (3x seeds vs. 2x seeds) > 1, and p-value < 0.05. This list was used to determine enriched GO-terms. Significantly enriched biological processes were identified with AtCOECIS (45) and further summarized using REVIGO (46).

To assess the individual behavior of auxin-related genes in 3x seeds, genes involved in biosynthesis, signaling, and transport of auxin were selected among the gene expression data produced by Schatlowski et al. (44), and their Log2FC (3x seeds vs. 2x seeds) values were plotted. Endosperm-specific expression of these genes was assessed using the transcriptome data of isolated endosperm from 3x and 2x seeds (47).

To evaluate expression changes of auxin-related genes throughout endosperm development, we used published transcriptome data (48). Only auxin-related genes that were expressed at the pre-globular stage, in a given endosperm domain, were considered for further analysis. Gene expression values in each endosperm domain and for each time point were then normalized to the pre-globular stage, and subsequently log transformed. To determine how the expression of these genes is affected in 3x seeds, where endosperm cellularization is restored, we used previously published transcriptome data of seeds corresponding to the cross L*er* x *osd1 adm-2* (13).

### Cloning and generation of transgenic plants

To clone the promoter of *DD25* (22, 26), WT Col-0 genomic DNA was used as a template. The amplified fragment was purified from the gel, recombined into the donor vector pDONR221 and sequenced. The insert was excised using the restriction sites SacI and SpeI introduced in the primer adaptors and used to replace the CaMV35 promoter in the vector pB7WG2 (49). The *IaaH* coding sequence was then recombined from an entry vector into pB7WG2, downstream of the *DD25* promoter, forming the *DD25::IaaH* construct. Gateway cloning was done according to the manufacturer’s instructions (Invitrogen). All primer sequences can be found in Table 2.

The construct was transformed into *Agrobacterium tumefaciens* strain GV3101 and *Arabidopsis* plants were transformed using the floral dip method (50). Transformants were selected with the appropriate antibiotics.

### Histological and fluorescence analyses

For clearing of ovules and seeds the whole pistils/siliques were fixed with EtOH:acetic acid (9:1), washed for 10 min in 90% EtOH, 10 min in 70% EtOH and cleared over-night in chloralhydrate solution (66.7% chloralhydrate (w/w), 8.3% glycerol (w/w)). The ovules/seeds were observed under differential interference contrast (DIC) optics using a Zeiss Axioplan or Axioscope A1 microscopes. Images were recorded using a Leica DFC295 camera with a 0.63x optical adapter.

For fluorescence analysis seeds were mounted in 7% glucose. Where indicated, 0.1 mg/mL propidium iodide (PI) was used. Samples were analyzed under confocal microscopy on a Zeiss 780 Inverted Axio Observer with a supersensitive GaASp detector with the following settings (in nm; excitation-ex and emission-em): GFP – ex 488, em 499-525; PI – ex 488/514, em 635-719; YFP (VENUS) – ex 514, em 499–552 for DR5v2. Images were acquired, analyzed and exported using Zeiss ZEN software.

For Feulgen staining of seeds, whole siliques were fixed in ethanol:acetic acid (3:1) overnight. The samples were washed three times 15 min in water, followed by 1 h incubation in freshly prepared 5 N HCl, and washed again three times 15 min in water. Staining was performed for 4 h in Schiff reagent, followed by three 15 min washes in cold water and a series of 10 min washes in a series of ethanol dilutions (10, 30 and 50%). The samples were then incubated in 70% ethanol overnight, which was followed by a 10 min wash in 95% ethanol and 1 h in 99.5% ethanol. Embedding of the seeds was done in a dilution series of ethanol:LR White resin (1:3, 1:2, 1:1, 2:1) for 1 h each. The samples were then incubated overnight in LR White resin, mounted in LR White plus accelerator and baked overnight at 60°C for polymerization. The seeds were imaged in a Zeiss multiphoton LSM 710 NLO with excitation at 800 nm and emission between 565–610 nm. The images were treated using the ZEN software.

### RT-qPCR analyses

For the determination of gene expression of *PEGs* and *AGLs*, ten whole siliques were collected for each cross and frozen in liquid nitrogen. All samples were collected in duplicate. Total RNA was extracted using the MagJET Plant RNA Purification Kit (Thermo Fisher Scientific) and 200 ng of total RNA were used to synthesize cDNA using the RevertAid First Strand cDNA Synthesis Kit (Thermo Fisher Scientific) using an oligo dT primer. Maxima SYBR Green qPCR Master Mix (Thermo Fisher Scientific) was used to perform the qPCR in a CFX Connect Sytem (Bio-Rad). The primers used for the RT-qPCR are described in Table S1. *PP2A* was used as the reference gene. Relative quantification of gene expression was performed as described (51).

## Acknowledgements

We are indebted to Dolf Weijers for providing the *DR5v2* reporter prior to publication, and to Eva Sundberg and Izabela Cierlik for providing the *IaaH* pENTRY vector.

---

**Figure 1-S1.**
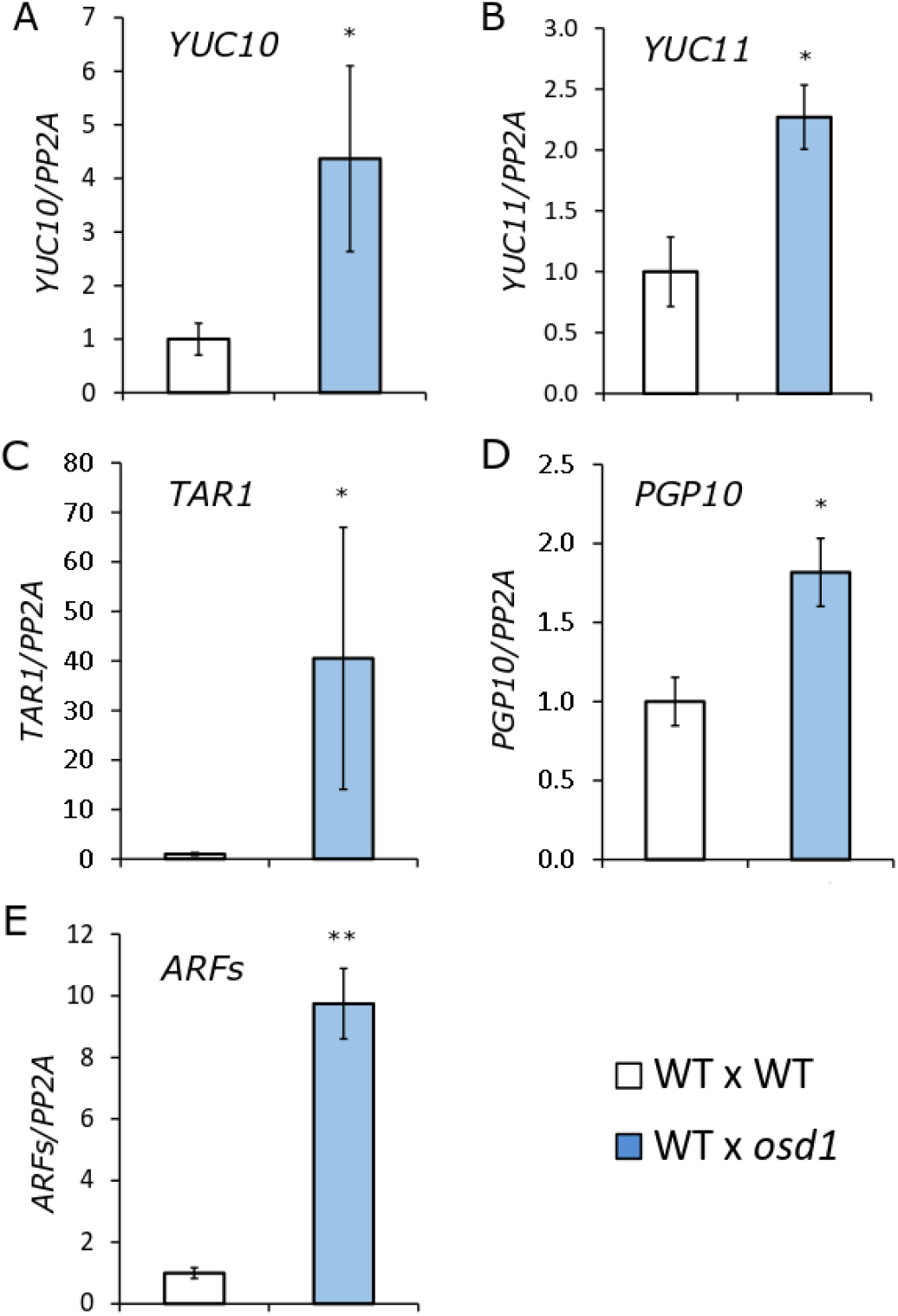
Validation of auxin-related gene expression from Fig. 1A. Relative gene expression in 2x and 3x seeds 6 days after pollination (6 DAP), as determined by RT-qPCR for *YUC10* (A), *YUC11* (B), *TAR1* (C), *PGP10* (D) and *ARFs* (E). Each figure panel shows a representative biological replicate. *ARF12, 13, 14, 15, 21* and *22* were assayed together due to high sequence similarity. Three technical replicates were performed and error bars indicate standard deviation. Differences are significant for Student’s T-test for p<0.05 (*) or p<0.001 (**).

**Figure 1-S2.**
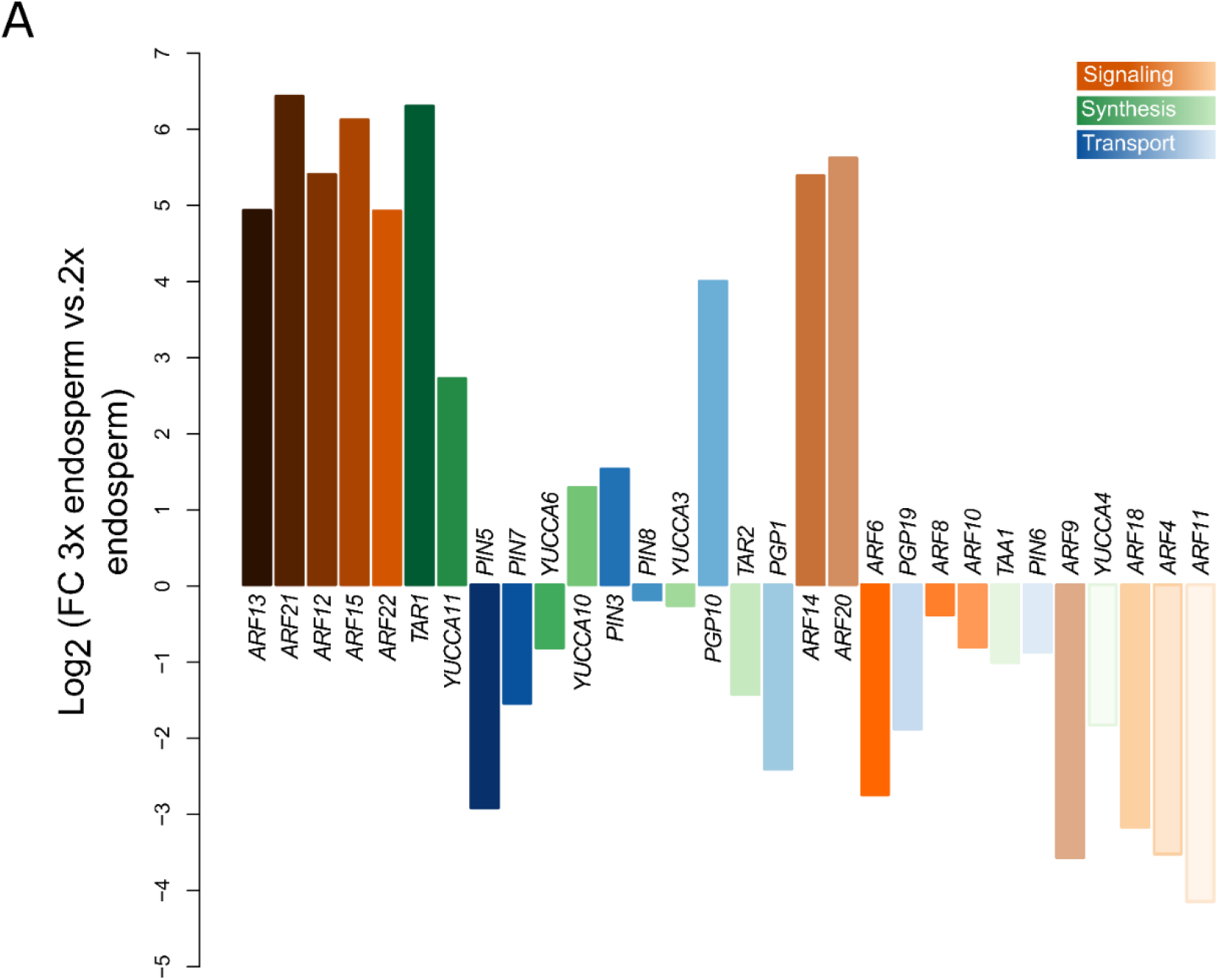
Auxin-related genes are upregulated in the endosperm of 3x seeds. Log2-fold expression change between genes expressed in the endosperm of 3x and 2x seeds. Genes coding for auxin signaling, biosynthesis, and transport proteins are indicated by orange, green and blue colors, respectively.

**Figure 2-S1.**
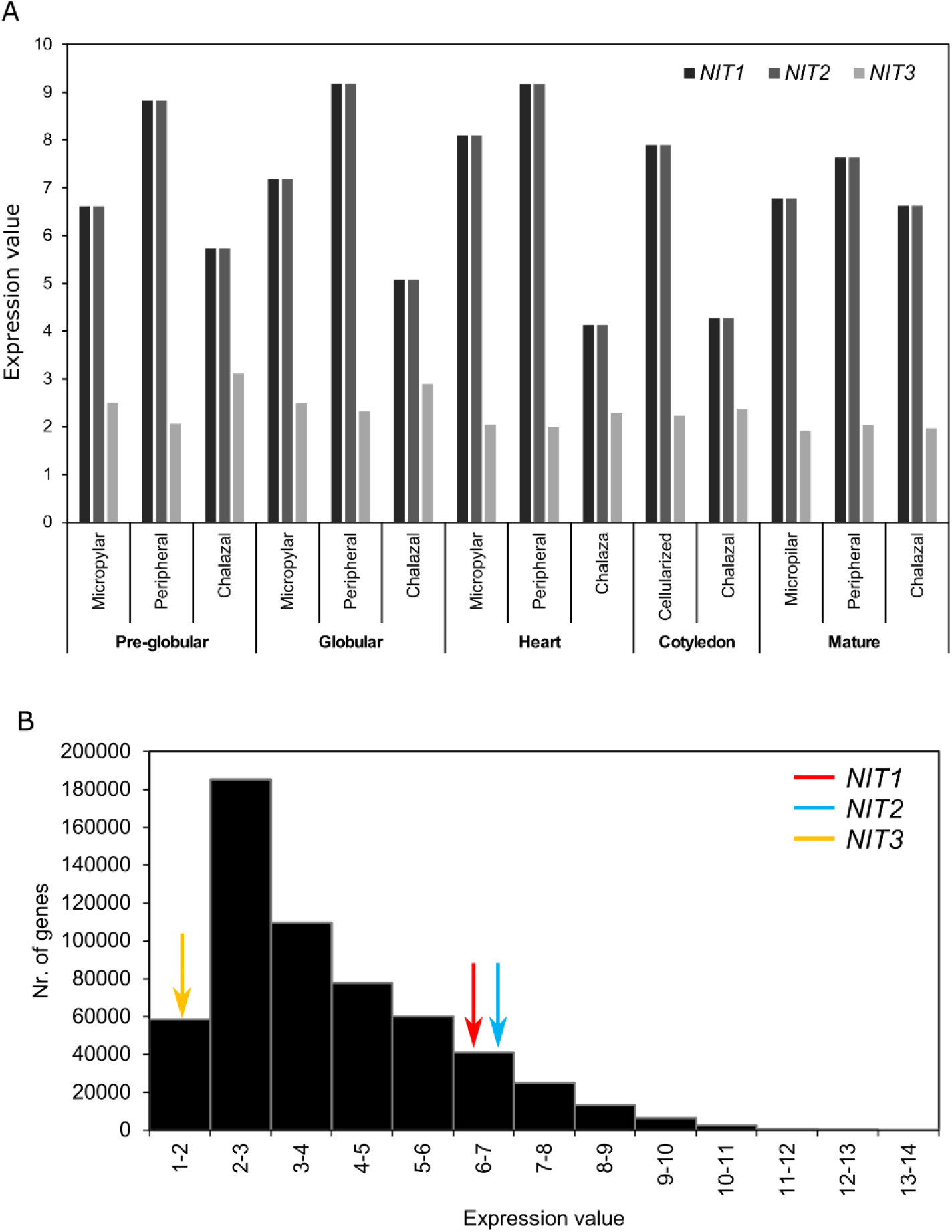
Relative expression of NITRILASE (NIT) coding genes. (A) Expression of genes coding for NIT IAM-synthetizing enzymes in each endosperm domain along seed development. Gene expression values indicate normalized microarray signal intensity according to (48). (B) Relative expression of *NIT1-3* genes, compared to the global distribution of gene expression in different subdomains and time-points of endosperm development, according to (48). Arrows indicate the gene expression category for each *NIT* gene.

**Figure 2-S2.**
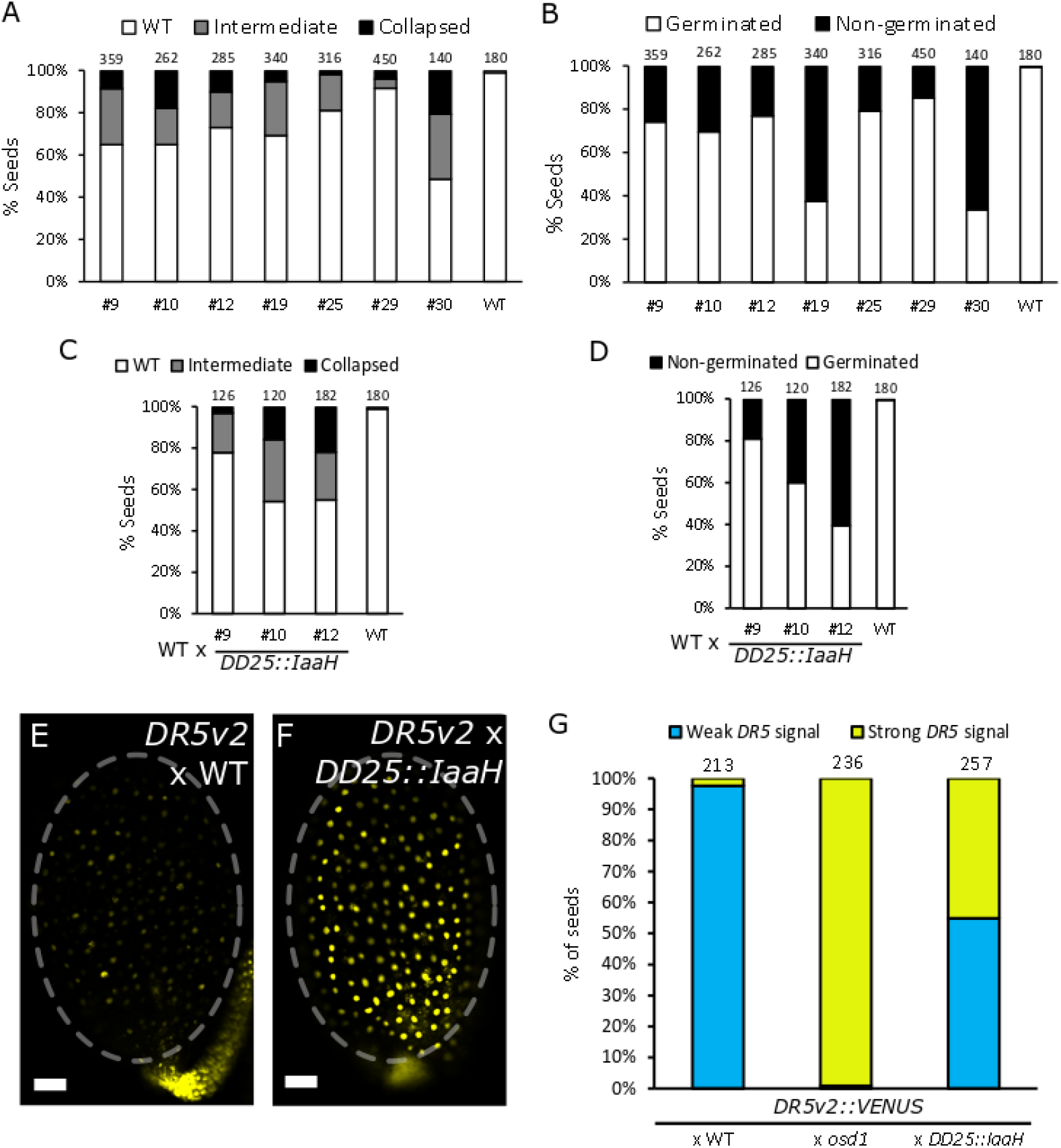
Expression of *DD25::IaaH* induces seed abortion. (A) Quantification of seed phenotypes in independent lines expressing *DD25::IaaH*. The seeds were classified in three distinct classes, as shown in Fig. 2-S3. (B) Seed germination rates in lines expressing *DD25::IaaH*. (C, D) Same as for (A) and (B), but using *DD25::IaaH* as pollen donor crossed to WT. Numbers on top indicate number of seeds analyzed. (E-F) Activity of maternal *DR5v2* at 5 days after pollination with WT (E) or *DD25::IaaH* (F) pollen. Pictures show representative seeds of three independent siliques per cross. Scale bars indicate 50 μm. (G) Percentage of seeds showing increased *DR5v2::VENUS* expression after pollination with a WT, *osd1* or DD25::IaaH-expressing father. For *DD25::IaaH*, line #10 was used. The seeds were categorized as showing weak or strong *DR5* signals according to panels (E) and (F), respectively. WT, wild type.

**Figure 2-S3.**
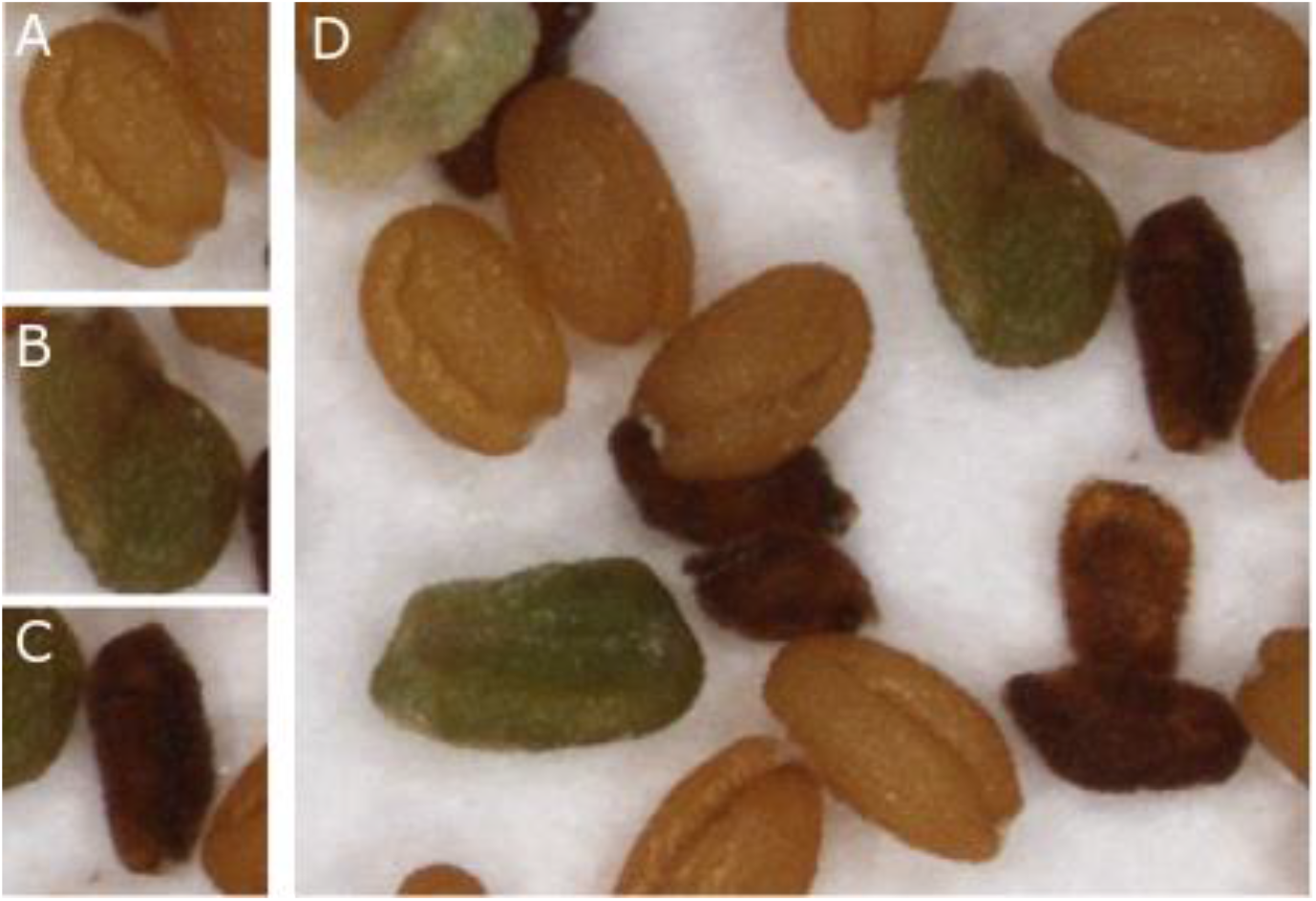
Seed classification used for the phenotypic quantification of *DD25::IaaH* phenotypes in Fig. 2-S2 and Fig. 4. (A) WT-like seed, (B) Intermediate phenotype, misshapen seeds, (C) Fully collapsed and shriveled seed. (D) Overall view of the progeny of a *DD25::IaaH* transgenic line.

**Figure 2-S4.**
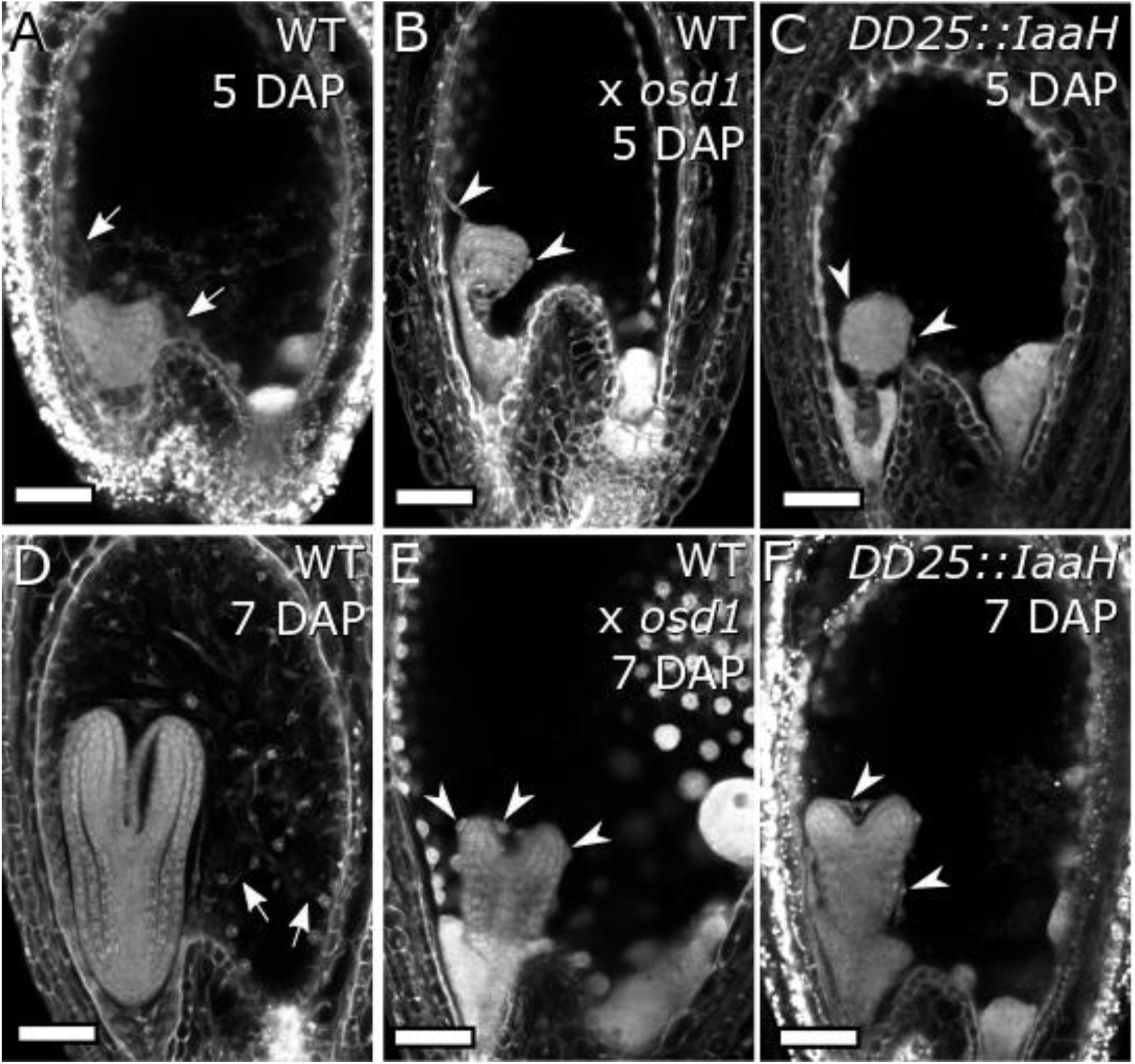
Endosperm cellularization as determined by Feulgen staining. (A-C) Seeds at 5 days after pollination (DAP) of WT 2x (A), WT 3x (B) and *DD25::IaaH* 2x (C). (D-F) Same as for (A-C), but for 7 DAP seeds. Pictures show representative seeds of 10 independent siliques per cross. Arrows indicate cellularized endosperm and arrowheads indicate free endosperm nuclei surrounding the embryo. Scale bars indicate 50 μm. WT, wild type.

**Figure 4 – S1.**
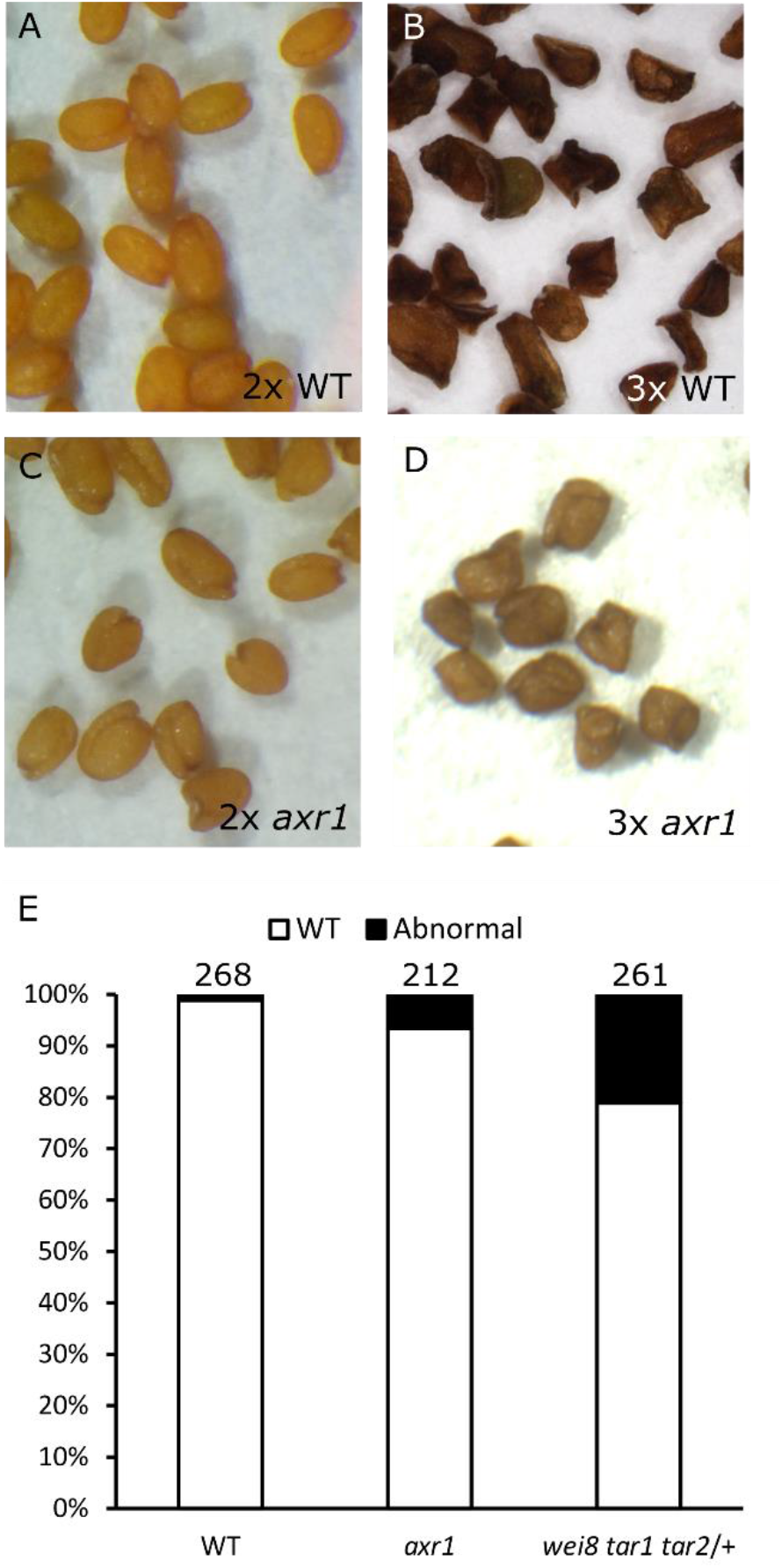
Seed phenotypes of 2x and 3x seeds. Mature seeds of WT Col-0, 2x (A) and 3x (B), and *axr1* mutant, 2x (C) and 3x (D). (E) Quantification of seed malformation in 2x seeds of wild-type (WT), *axr1* and *wei8 tar1 tar2/+* plants. Numbers on top indicate number of seeds assayed.

**Figure 4 – S2.**
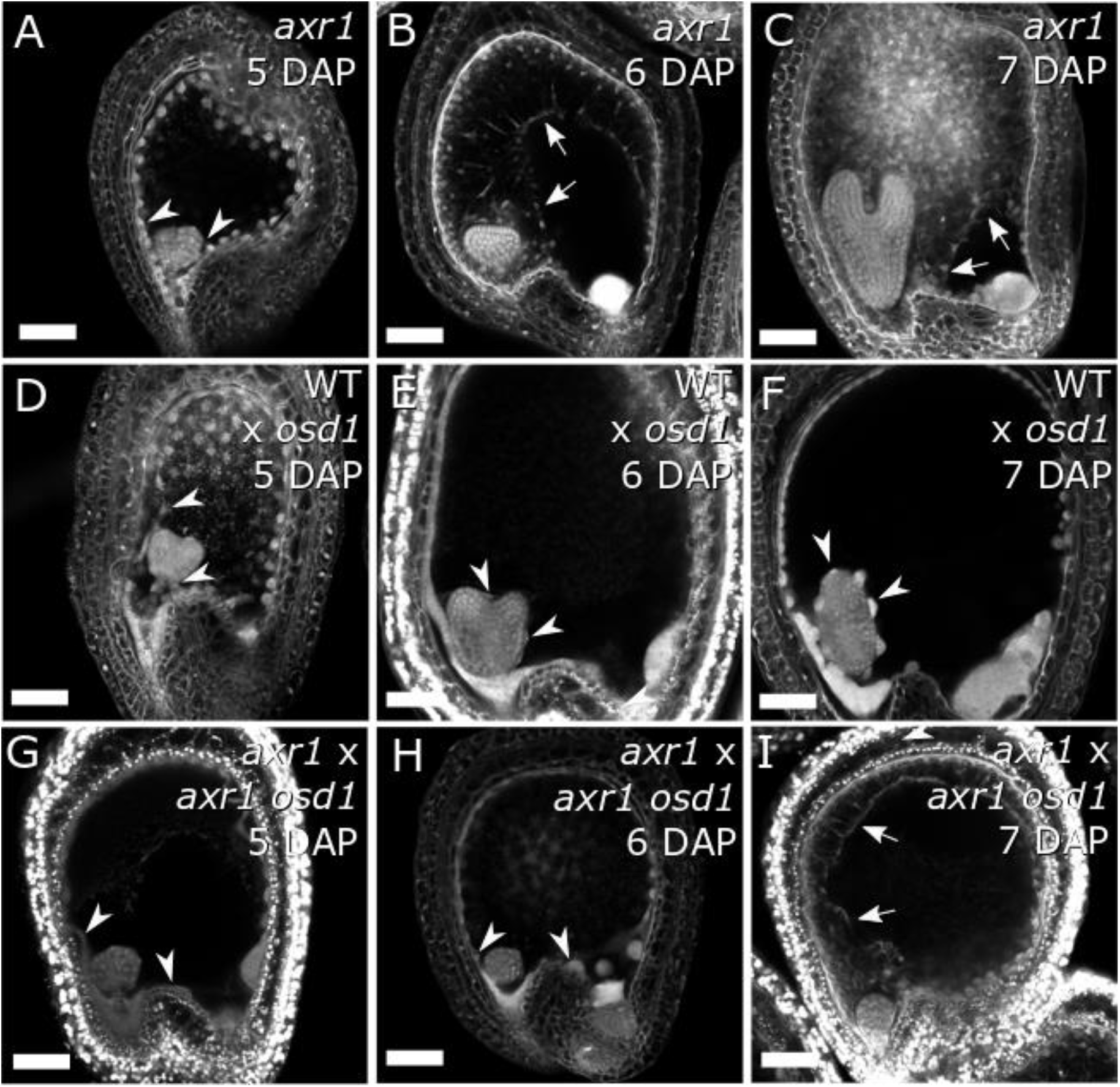
Loss of *AXR1* function restores endosperm cellularization in 3x seeds. A-C) 2x *axr1* seeds at 5, 6 and 7 days after pollination (DAP). (D-F) Same as for (A-C), but for 3x WT seeds. (G-I) Same as (A-C), but for 3x *axr1* seeds. Pictures show representative seeds of 10 independent siliques per cross. Arrows indicate cellularized endosperm and arrowheads indicate free endosperm nuclei surrounding the embryo. Scale bars indicate 50 μm. WT, wild type.

**Figure 5-S1.**
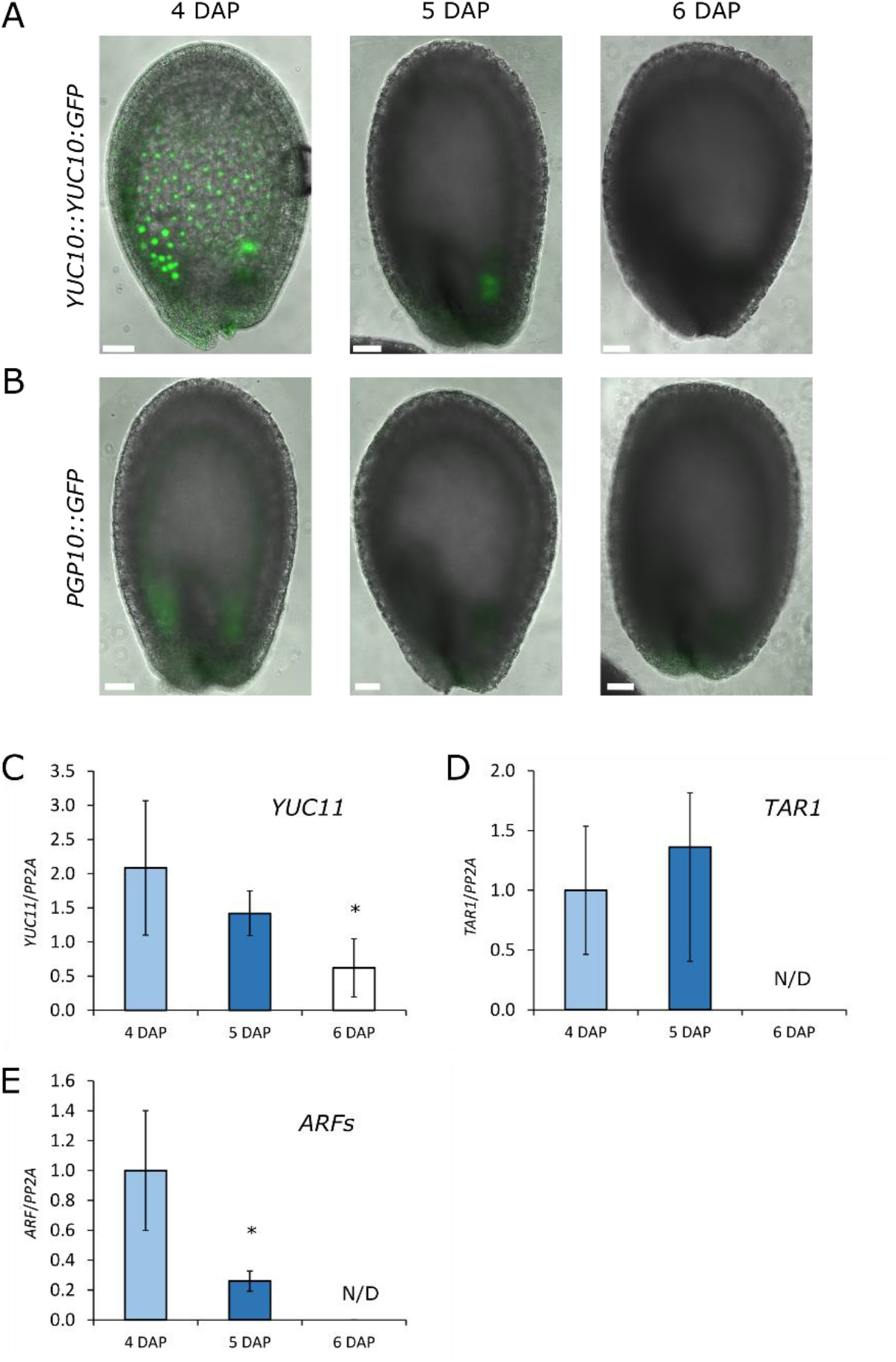
Downregulation of auxin-related genes coincides with the timing of endosperm cellularization. (A-B) Reporter activity for *YUC10::YUC10:GFP* (A) and *PGP10::GFP* (B) during seed development (4 to 6 days after pollination, DAP). Scale bars indicate 50 μm. (C-E) Relative gene expression in seeds at 4, 5 and 6 DAP, as determined by RT-qPCR for *YUC11* (C), *TAR1* (D), and *ARFs* (E). Each figure panel shows a representative biological replicate. *ARF12,13,14,15, 21* and *22* were assayed together due to high sequence similarity. Three technical replicates were performed and error bars indicate standard deviation. Differences are significant for Student’s T-test for p<0.05 (*). N/D, transcript not detected.

**Table S1.**
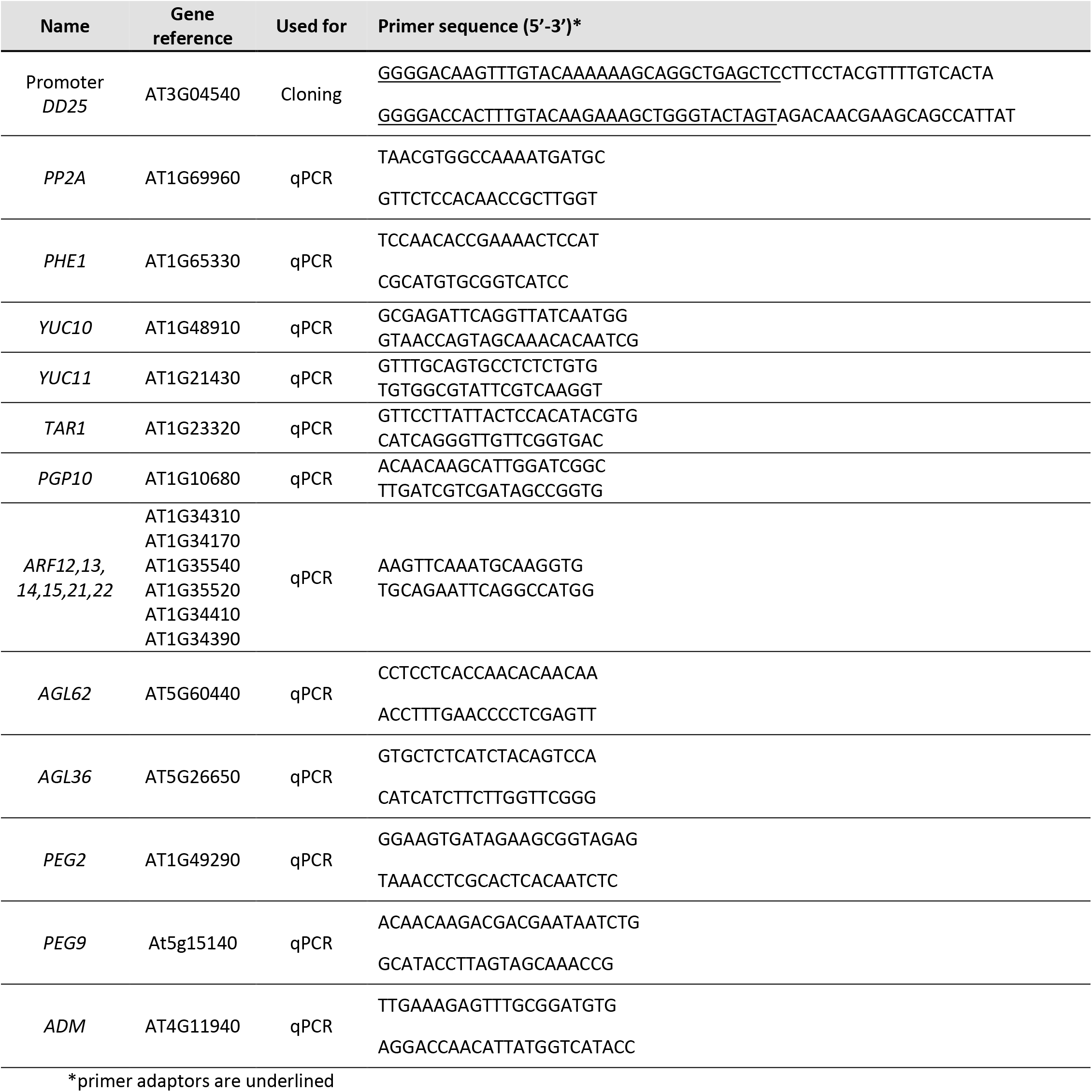
Primer list

